# Solving the transcription start site identification problem with ADAPT-CAGE: a Machine Learning algorithm for analysis of CAGE data

**DOI:** 10.1101/752253

**Authors:** Georgios K Georgakilas, Nikos Perdikopanis, Artemis Hatzigeorgiou

## Abstract

Cap Analysis of Gene Expression (CAGE) experimental protocol has emerged as a powerful experimental technique for assisting in the identification of transcription start sites (TSSs). There is strong evidence that CAGE also identifies capping sites along various other locations of transcribed loci such as splicing byproducts, alternative isoforms and capped molecules overlapping introns and exons. We present ADAPT-CAGE, a Machine Learning framework which is trained to distinguish between CAGE signal derived from TSSs and transcriptional noise. ADAPT-CAGE provides annotation-agnostic, highly accurate and single-nucleotide resolution experimentally derived TSSs on a genome-wide scale. It has been specifically designed aiming for flexibility and ease-of-use by only requiring aligned CAGE data and the underlying genomic sequence. When compared to existing algorithms, ADAPT-CAGE exhibits improved performance on every benchmark that we designed based on both annotation- and experimentally-driven strategies. This performance boost brings ADAPT-CAGE in the spotlight as a computational framework that is able to assist in the refinement of gene regulatory networks, the incorporation of accurate information of gene expression regulators and alternative promoter usage in both physiological and pathological conditions.

## Introduction

Cap Analysis of Gene Expression (CAGE) was initially introduced in 2003 [1] as a novel method specifically developed to capture and quantify 5’ ends of capped RNAs. During the last decade, CAGE has been continuously refined and improved into its current mature form as a well-established protocol for the identification of transcription start sites (TSS) and promoter regions of transcribed loci. The FANTOM Consortium [2] has extensively applied CAGE on hundreds of tissues and cell-lines to produce a high-quality annotation of the human and mouse promoterome and characterize regulatory mechanisms of gene expression.

Despite its increasing popularity as an experimental promoter identification protocol, the specificity of CAGE regarding the identification of transcription initiation events in the genome has several limitations. There is strong evidence [3–5] that besides promoter regions, CAGE also identifies capping sites along various locations of transcribed loci such as different splicing products, isoforms and capped molecules that can be summed up as transcriptional noise. As a result, only a portion of regions enriched in CAGE signal were found to overlap with the surrounding region of annotated TSSs.

This poses a significant hindrance to research studies that aim to integrate regulatory regions into the framework of biological pathways. During the last decade, several *in silico* methodologies have been developed to provide basic pipelines for analyzing CAGE datasets and to facilitate peak identification and annotation. Paraclu [6] performs clustering of CAGE tags into wider regions based on a density parameter that reflects the trade-off between size and number of overlapping reads. Within clusters, a score that represents the likelihood of being a transcription initiation site is derived from the analysis of surrounding genomic sequence using a position-specific Markov Model trained on k-mer frequencies. RECLU [7] is an adaptation of Paraclu algorithm that is able to handle replicated (max two replicates) experiments based on the irreproducible discovery rate (IDR) algorithm [8] and utilizes slightly modified parameters to filter the final results when compared against the original implementation. CAGEr is the most recent *in silico* framework for CAGE analysis and TSS identification [9]. CAGEr applies quality filtering, and depending on the utilized protocol, removes the 5′ end G nucleotide bias. It subsequently clusters reads into groups and is able to handle multiple CAGE experiments, detect differential TSS usage while addressing the in-between tissue variability in TSS choice and promoter shifting.

Despite the important advances in the development of CAGE analysis pipelines, it is evident that existing implementations still include a high number of false positive rate on TSS identification in CAGE datasets. In various older studies, unique structural features were found to be associated with promoter regions [10–13]. In a recent study [10], the structural properties of DNA in promoter regions were investigated by analyzing 13 features including duplex disrupt energy, duplex free energy, bending stiffness, denaturation, stacking energy, bendability, propeller twist, z-DNA, A-philicity, nucleosome positioning, protein deformation, B-DNA twist and protein-DNA twist. Based on structural models derived from a wide array of biochemical experiments, sequences around transcription initiation events from DBTSS (positive set) were converted into numerical vectors and compared against their shuffled counterparts (negative set). It was shown that there are distinct patterns of structural DNA features that can distinguish between promoter and non-promoter regions. Based on these observations we hypothesized that the same features and their promoter patterns should be able to facilitate the classification of CAGE tag-clusters between real TSSs and transcriptional noise.

In this study, we present ADAPT-CAGE, a versatile computational framework for analyzing CAGE data providing annotation-agnostic, highly accurate and single-nucleotide resolution experimentally derived TSSs on a genome-wide scale. The proposed algorithm (Figure 1) introduces a novel approach for distinguishing CAGE tag-clusters that represent transcription initiation events from clusters that are formulated due to recapping events, byproducts of the splicing machinery as well as transcriptional and/or sequencing noise. Initially, CAGE tags are filtered based on their mapping quality and 5’ ends of the remaining reads are combined into clusters depending on their in-between distance. For each CAGE peak, the normalized number of overlapping reads is computed and the representative nucleotide is selected based on the localization of CAGE tag 5’ ends. The algorithm strategically utilizes sequence and structural DNA properties that have been previously [10] associated to transcription initiation and Polymerase II promoter regions. Based on the sequence surrounding each representative TSS, ADAPT-CAGE extracts sequence and structural DNA features which are then forwarded into a multilayered Machine Learning module. This module is based on Support Vector Machines (SVM) and Stochastic Gradient Boosting (SGB) models individually trained on each DNA feature and subsequently combined with an agent assembly strategy.

**Figure 1.**
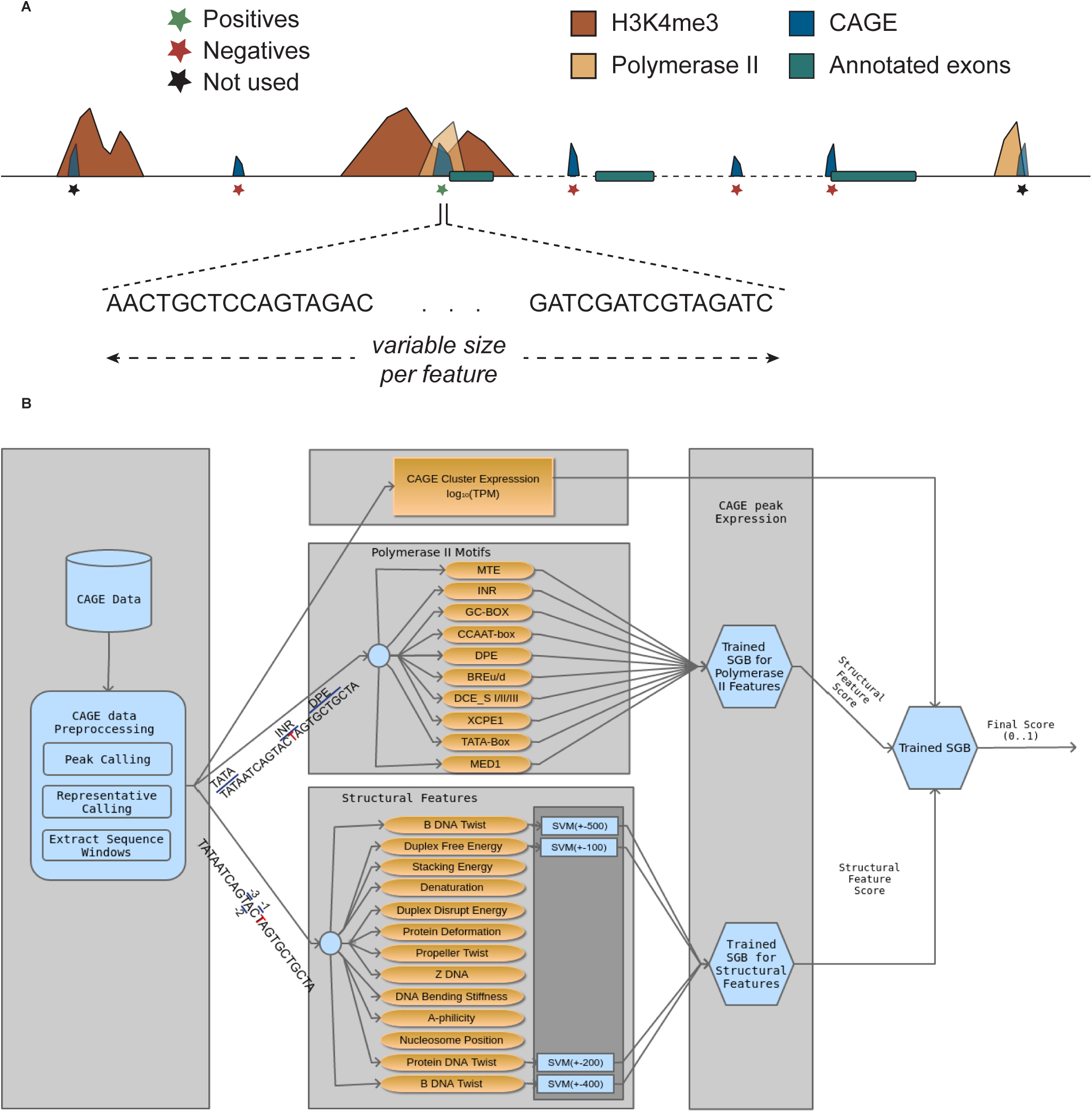
Overview of ADAPT-CAGE and the training process. A) CAGE clusters selected for the positive set originated from annotated promoter regions with H3K4me3 and Polymerase II ChIP-Seq derived occupancy. Negative samples were selected from intronic, exonic and intergenic regions that did not overlap H3K4me3 or Polymerase II peaks. B) ADAP-CAGE accepts as input aligned reads in BAM format. The preprocessing module aggregates CAGE reads into peaks and finds their representative nucleotide (the position exhibiting the highest expression within each peak). The sequence surrounding the representatives is extracted and the structural as well as Polymerase II features are calculated and forwarded on their own GBM (Polymerase II motifs) and SVM (structural features) models. The last module of the framework is based on a GBM model that combines previous outputs with the expression level of candidate TSSs to produce the final output of the algorithm.

## Results

### Genome-wide assessment of FANTOM CAGE tag cluster quality, pre- and post-ADAPT-CAGE application

There is increasing evidence in the literature [5,14] that unveils CAGE’s property to detect recapping events, alternative isoforms and in some cases, post-transcriptional processing and cleavage by RNA binding proteins, in addition to transcription initiation events. To showcase the aforementioned properties, we adopted an unsupervised exploratory strategy on a genome-wide scale utilizing ChIP-Seq data against H3K4me3, H3K4me1, H3K27ac, H3K27me3, H3K9me3 and H3K36me3. These histone marks are traditionally used to characterize chromatin states in landmark studies and projects such as Roadmap Epigenomics [15].

More than 31,000 CAGE enriched regions were identified in H9 ES cells (as reported in FANTOM repository) and the occupancy of each histone mark was calculated in their surrounding region (+/− 1kb). The histone mark enrichment formed the basis for clustering these regions into groups with similar patterns of chromatin activity. Normalized signal profiles were also added as a visual aid. In Figures 2A and 2B, two active loci (SEMA4C and TMEM131) are shown along with the normalized CAGE, ChIP- and DNase-Seq signal depicting different types of activity. In both cases, DNase-Seq and H3K4me3 ChIP-Seq data indicate that both promoters are accessible and CAGE shows that the two genes are expressed. We can observe numerous CAGE enriched regions in their exons and introns, especially in the case of TMEM131 whose length is above 200kb.

**Figure 2.**
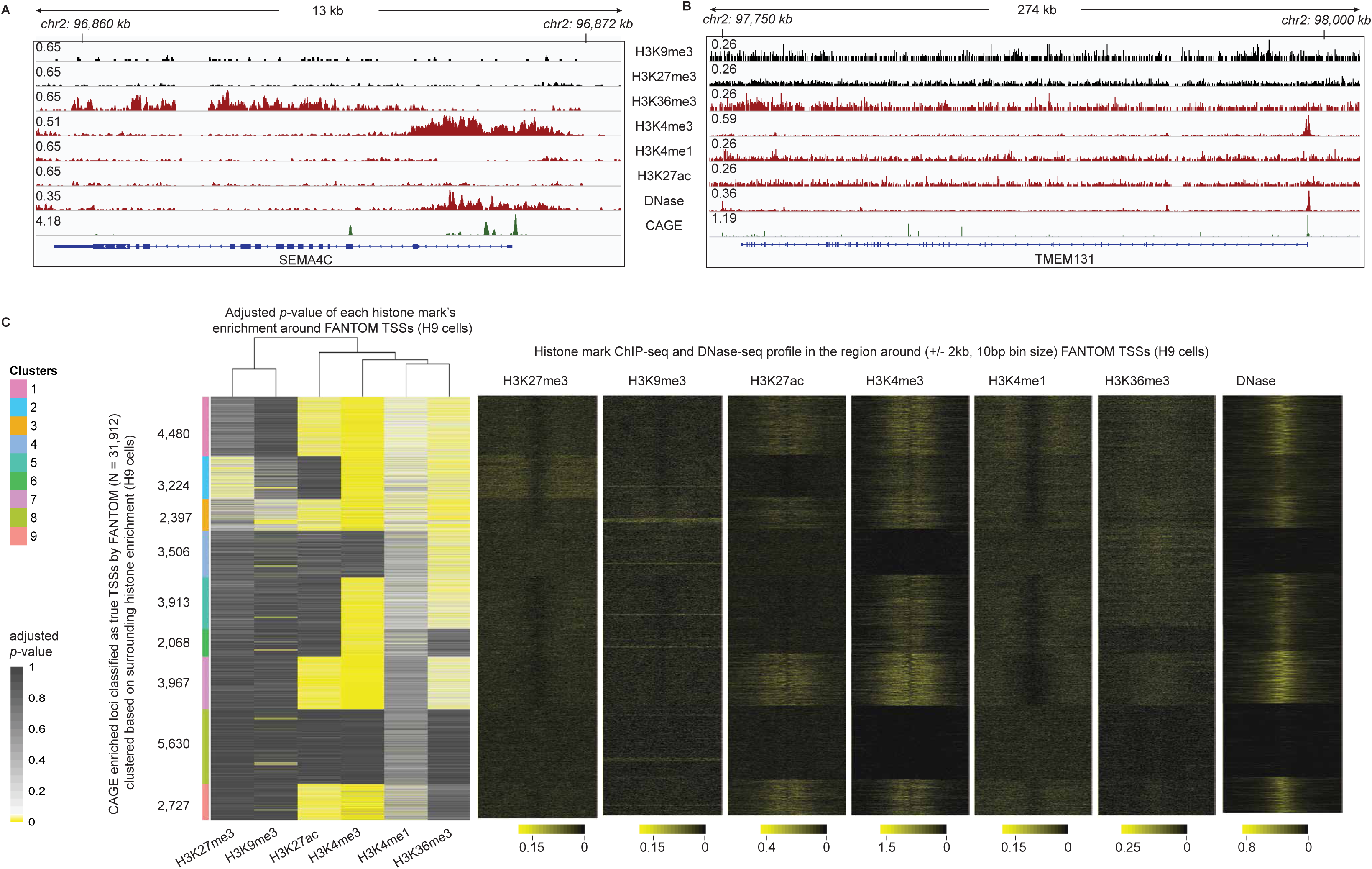
Genome-wide analysis of the chromatin landscape surrounding CAGE enriched loci in H9 cells as reported by FANTOM. Examples of A) SEMA4C and B) TMEM131 loci with tracks from six histone marks showing chromatin status, DNase-Seq unveiling accessibility and the actual CAGE signal. Active or poised promoters are denoted with H3K4me3 and H3K27ac, active or poised enhancers with H3K4me1 and H3K27ac, repressed chromatin with H3K27me3 and H3K9me3, and transcribed regions with H3K36me3. As indicated by the CAGE signal, there are multiple CAGE enriched regions located in introns and exons of SEMA4C and especially of TMEM131 with length over 200kb. The ones located in promoter regions exhibit increased chromatin accessibility while the rest are located in inaccessible chromatin. C) 31,912 FANTOM CAGE peaks were clustered based on the enrichment of six histone marks (H3K4me3, H3K27ac, H3K4me1, H3K36me3, H3K9me3, H3K27me3). On the right side, histone ChIP-, DNase-Seq and CAGE signal profile is also shown. We observe that 61.3% (19,552) CAGE peaks exhibit active histone mark enrichment and the remaining 38.7% (12,360) do not demonstrate any chromatin activity showcasing the increased transcriptional noise level embedded in CAGE datasets.

Similar activity patterns are also observed on the genome-wide scale (Figure 2C). There are two major chromatin modes around CAGE enriched loci. Most of them (61.3%, 19,552) exhibit, as expected, a medium to strong enrichment in DNase- and histone marks ChIP-Seq related to active promoters/enhancers. The remaining loci (38,7%, 12,360) in clusters 2, 4 and 8, do not exhibit enrichment in DNase-seq signal nor in any of the histone marks. These findings are in tandem with the evidence from the literature, highlighting the significant enrichment of CAGE datasets in signal that corresponds to transcriptional noise.

The question that arises is how the picture of the chromatin landscape around the aforementioned CAGE enriched regions changed after applying ADAPT-CAGE. To test this, we filtered these regions using an ADAPT-CAGE score of 0.5 resulting in 17,850 loci passing (Supplementary Figure 2A) and 14,062 below the cutoff (Supplementary Figure 2B). The vast majority (97.1%, 17,335 out of 17,850) of the positively scored CAGE enriched regions exhibit medium or strong enrichment in histone modifications signal of active transcription. Cluster 9 consisting of only 515 regions is the only group that is not enriched in such signal. In contrast, 9,624 out of 14,062 (clusters 2, 5, 7, 8, 9 and 10 in Supplementary Figure 2B) loci that failed to pass the ADAPT-CAGE score cutoff do not exhibit any enrichment in active transcription signal, while the remaining 4,438 presents medium to strong activity.

The evidence presented in this section support that ADAPT-CAGE is able to correctly distinguish between CAGE enriched regions associated to transcription initiation events from noise. However, even though ADAPT-CAGE exhibits very high levels of precision, the performance is terms of sensitivity seems to be lagging behind in this analysis. In general, every unsupervised learning strategy inherently produces results with increased within-cluster variability and k-means is not the optimal algorithm for dealing with non-linearity. In addition, the lower levels of sensitivity could also be explained by the limitations of the experimental methods used to produce the genome-wide histone occupancy and the inability to detect lowly expressed regions due to the selected sequencing depth. In any case, the unsupervised strategy presented here provides only a rough estimate of ADAPT-CAGE’s performance. In the following sections, we present more results from benchmarking strategies based on additional experimental evidence and the annotation of the human genome as well as performance comparisons with existing algorithms. The number of total predictions for each algorithm are listed in Supplementary Table 1.

### Experimentally- and ChromHMM-driven comparison between ADAPT-CAGE and existing algorithms

The Roadmap Epigenomics [15] and ENCODE [16] projects have long been a one-stop data repository for epigenomics and transcriptomics providing hundreds of high quality samples from multiple tissues and cell lines. This wealth of information enabled the development of numerous computational methods such as ChromHMM [17] which has emerged as a powerful approach for annotating the epigenome based on experimentally-driven identification of cell-specific chromatin states.

To further explore the limits of ADAPT-CAGE’s performance and compare it with existing algorithms, we developed a benchmarking strategy based on the core-15 [17,18] chromatin states from ChromHMM (Supplementary Table 2) in H9 and K562 cells. Initially, for every algorithm we calculated the percentage of predictions that overlap each chromatin state (Figure 3A,B and Supplementary Figure 3A,B). The chromatin states were aggregated into two groups based on their relation with active or repressed chromatin. In the active chromatin group (TssA, TssAFlnk, TxFlnk), the top-performing algorithm is ADAPT-CAGE (94.83%) followed by PARACLU (89.84%), RECLU (84.82%), and CAGER (84.31%). ADAPT-CAGE also exhibits the lowest percentage in the repressed chromatin group (5.16%), followed by PARACLU (10.15%), RECLU (15.17%) and CAGER (15.68%). The results for the K562 comparison are shown in Supplementary Figure 3A,B.

**Figure 3.**
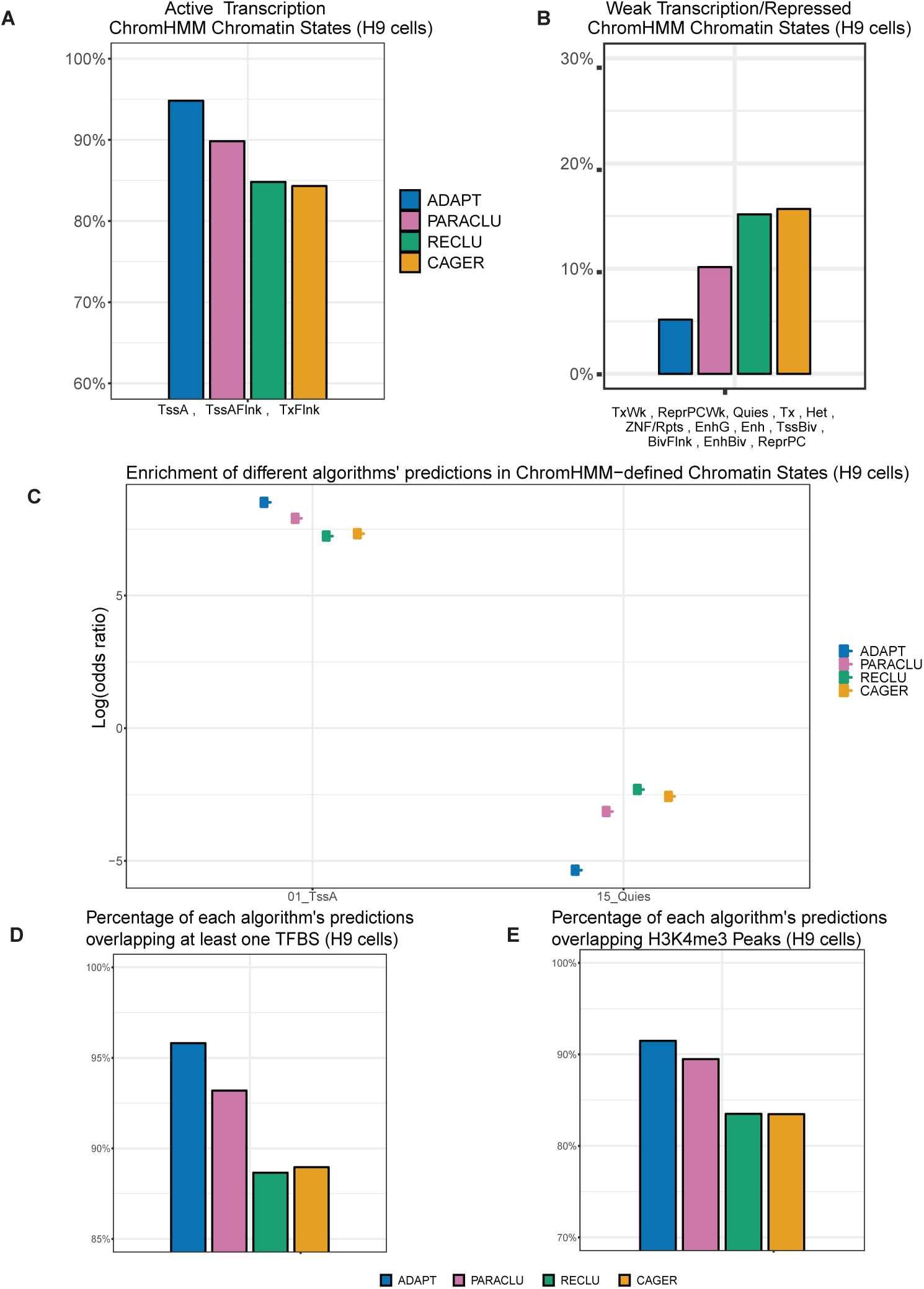
Evaluating algorithms’ performance on experimental data in H9 cells. Percentage of each algorithm’s predictions uniquely overlapping ChromHMM-derived chromatin states based on the core-15 model in H9 cells. Chromatin states were aggregated in two groups according to levels of chromatin/transcriptional activity. Active (A) and weaker (B) transcription states. C) Enrichment (odds ratio in logarithmic scale) of algorithms’ prediction in the two most orthogonal ChromHMM-derived chromatin states (TssA and Quies). Percentage of each algorithm’s predictions overlapping at least one transcription factor binding site (D) and H3K4me3 (E) ChIP-Seq derived peaks.

To compare the performance in a more unbiased way that alleviates the different number of predictions provided by each algorithm, we proceeded with the calculation of the odds ratio (OR) for the enrichment of positively scored predictions in each chromatin state, in logarithmic scale (Figure 3C). The OR for each algorithm in the two most orthogonal, in terms of activity, states is shown. TssA state characterises active TSS-proximal promoters and Quies state describes regions with no histone enrichment. In both cases ADAPT-CAGE outperforms all other algorithms. The results for the K562 comparison are shown in Supplementary Figure 3C.

To further explore the algorithms’ performance without using any third-party computational method, transcription factor binding sites (TFBS) were downloaded from the ENCODE’s Txn factor track that incorporates ChIP-Seq derived binding sites for more than 160 transcription factors. The overlap of each algorithm’s prediction with at least one TFBS in H9 cell lines was calculated (Figure 3D). ADAPT-CAGE achieves the best performance (95.81%) followed by PARACLU (93.19%), RECLU (88.65%) and CAGEr (86.96%). Similar results are shown in Supplementary Figure 3D for the same comparison based on K562 data.

The last benchmark in the experimentally oriented group of comparisons was based on peaks derived from ChIP-Seq data against H3K4me3, a histone modification that is highly enriched at active promoters (Figure 3E). CAGE enriched regions that were positively scored by ADAPT-CAGE demonstrate the highest overlap with H3K4me3 peaks (91.48%), followed by PARACLU (89.48%), RECLU (83.49%), and CAGEr (83.47%). Similar observations were made in the comparison based on K562 data (Supplementary Figure 3E).

### Annotation-driven comparison between ADAPT-CAGE and existing algorithms

For a further systematic evaluation of algorithms that can be applied on CAGE enriched regions we used the RefSeq protein coding gene annotation [18,19]. The region surrounding annotated TSSs (+/− 50kb) were selected to form the set of benchmark parts of the genome (see Materials and Methods). Initially, all CAGE peaks located +/− 500bp around TSSs were considered as positive, while the ones located in the remaining parts of the benchmark regions were treated as negative (Figure 4A). The same strategy has previously been used in [2]. In the second approach, we expanded the positive zone to +/− 1kb. This definition of positive and negative zones enabled the calculation of performance metrics such as specificity and sensitivity in H9 cell line.

**Figure 4.**
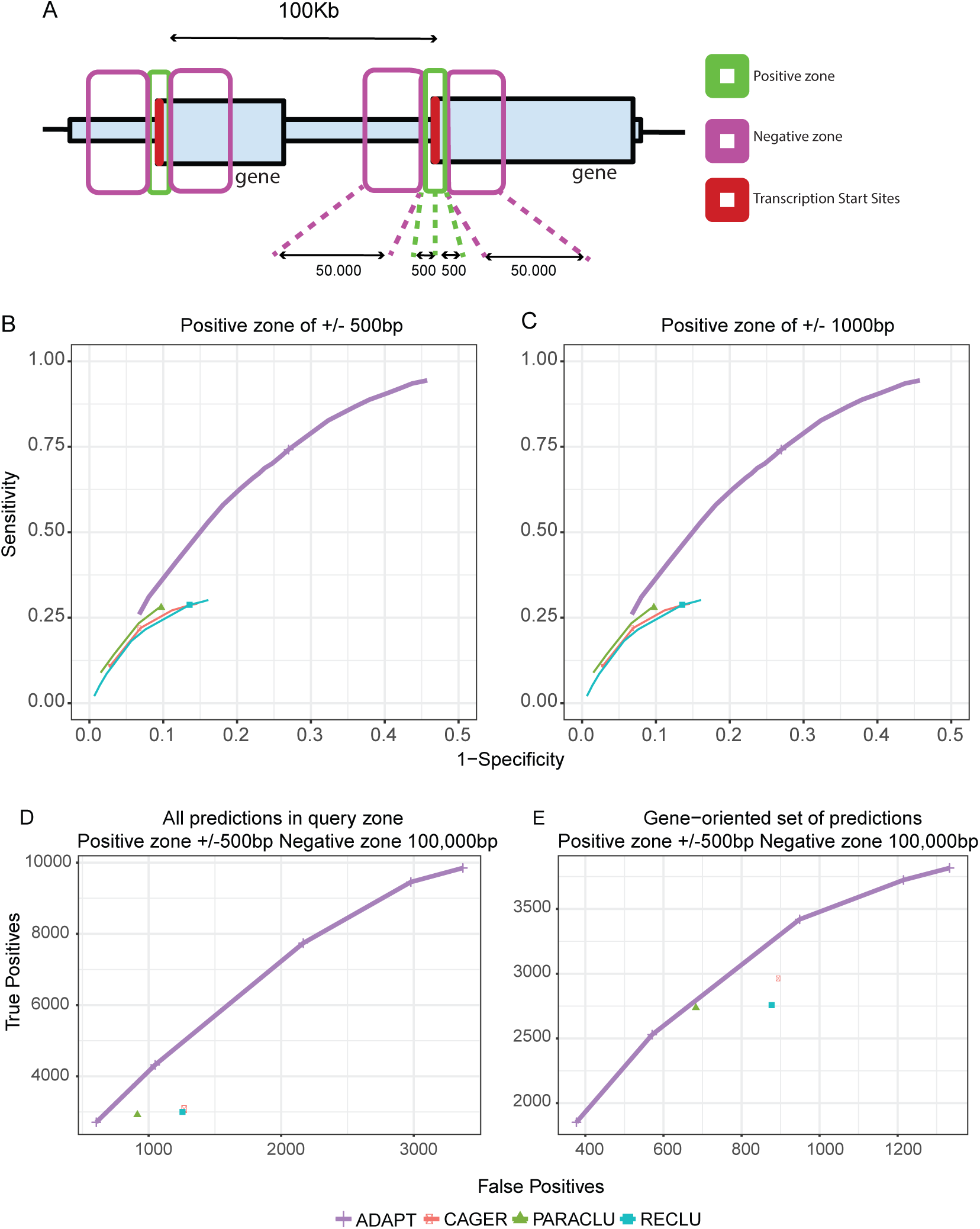
Evaluating algorithms’ performance on the human genome annotation in H9 cells. For a more systematic comparison of the algorithms’ performance, the human genome annotation was utilized for generating positive and negative zones surrounding annotated protein coding TSSs (A). The positive zone was defined as a +/− 500bp window centered on annotated TSSs. Predictions in this zone were accepted as true positives. In the remaining region outside this zone and up to +/− 50kb away from TSSs (negative zone), all predictions were accepted as false positives. See Materials and Methods for more details on the selection on positive and negative zones. ROC curves based on two different positive zone sizes, +/− 500bp (B) and +/− 1kb (C). The curves were generated by applying multiple ADAPT-CAGE score cutoffs and a range of different parameters for the application of existing algorithms (from loose to stricter outcomes). The complete range of utilized parameters are shown in Supplementary Table 4. Number of true positives plotted against false positives based on the +/− 500bp positive zone (D), and based on a gene-oriented strategy (E). In (E), a true positive prediction is a gene that has at least one prediction in the positive zone (+/− 500bp around its TSS) and a false positive prediction is a gene with at least one prediction in the negative zone.

In contrast to existing algorithms, ADAPT-CAGE is the only method that provides a score to every processed CAGE enriched region. Therefore, by applying multiple score cutoffs, we evaluated ADAPT-CAGE’s performance at each threshold. To perform a similar evaluation on existing algorithms we applied them using different parameters at each run, thus creating a set of different results (from loose to stricter outcomes). Using the +/− 500bp positive zone strategy, existing algorithms (Supplementary Table 2) have a comparable specificity to ADAPT-CAGE but they exhibit only a small range of sensitivity (Figure 4B). The same pattern is observed after increasing the positive zone to +/− 1kb (Figure 4C). In Figure 4D we present the results based on the +/− 500bp positive zone in absolute numbers of true positives (TPs) and false positives (FPs).

The previous benchmark was a CAGE peak centric approach. For the following comparison we adopted a gene oriented strategy. A positive prediction is a gene that has at least one hit in the +/− 500bp region around its TSS. The results are shown in Figure 4E depicting that ADAPT-CAGE outperforms all existing algorithms (Supplementary Table 1).

As a final comparison of algorithms in H9 cells, we counted the overlap of positively scored CAGE enriched loci with promoters, introns, exons and splicing sites (Supplementary Figure 5A). ADAPT-CAGE exhibits the highest overlap with promoters (92.7%), followed by PARACLU (88.36%), RECLU (83.12%) and CAGEr (82.47%). Additionally, ADAPT-CAGE presents the lowest overlap with other parts of genic loci (Supplementary Figure 5B).

## Discussion

Eukaryotic organisms exhibit diverse gene expression patterns, a feature that is responsible for the spatio-temporal cell diversity in such multicellular species. Unlocking the mechanisms that regulate transcription is a fundamental step towards understanding the rules of biological systems that when disrupted, diseases emerge. To achieve this, easy-to-implement experimental protocols and robust computational solutions are imperative. With the advent of Next Generation Sequencing, experimental methods able to quantify gene expression and/or transcript abundance have emerged. CAGE is one of these approaches.

CAGE was initially introduced in 2006 [1,15] and since then it has been continuously refined and improved into its current, significantly more mature, status. Nowadays, CAGE is considered a state-of-the-art protocol for experimental identification of transcription initiation events. Despite its popularity, many studies [5,14] have unveiled that CAGE is able to identify a wide array of biological events besides transcription initiation such as byproducts of the splicing machinery and transcriptional noise in general. Naturally, these observations suggest a significant number of false positive TSSs identified in CAGE datasets. This poses a significant obstacle to transcription-related research, signifying the need to develop computational methods for excluding events captured with CAGE that do not associate with TSSs.

In this study, we unveiled this problem on a genome-wide scale using an unbiased approach that involved unsupervised Machine Learning and epigenetic datasets (Figure 2). More than one third of the H9 CAGE tag-clusters provided in FANTOM database are located in repressed chromatin exhibiting zero transcriptional activity. After applying ADAPT-CAGE, we observed a significant improvement in identifying true transcriptional initiation events (Supplementary Figure 2). The limits of ADAPT-CAGE and existing algorithms’ performance was further explored using third-party algorithms such as ChromHMM, the human genome annotation and epigenomic data. Several observations were made regarding existing algorithms. Specifically, all algorithms involve stringent cutoffs in their pipeline resulting in the removal of valuable information from the CAGE tag-cluster set. The remaining tag-clusters exhibit high coverage in CAGE reads but that does not imply they are true positive transcription initiation events. On the other hand, ADAPT-CAGE is far more sensitive and harnesses the power of Machine Learning on promoter-associated and structural DNA features (Figure 1) to provide highly accurate predictions without removing the majority of candidate TSSs (Supplementary Table 1). It is able to perfectly complement research oriented around transcriptional mechanics by removing CAGE tag-clusters or genomic loci that are not associated with true TSSs.

In light of Next Generation Sequencing, accurate computational methods such as ADAPT-CAGE can emerge as key components of studies that aim to identify the regulatory machinery and core constituents of biological pathways. This is a fundamental endeavor towards understanding and characterizing the underlying mechanisms that govern every aspect of physiological conditions in biological systems, as well as stimuli that cause the transition into pathological states.

## Materials and Methods

### CAGE- and ChIP-Seq data analysis and utilized annotation

For the training and evaluation process we used pre-processed CAGE reads aligned on GRCh38 assembly of the human genome. The training of ADAPT-CAGE was achieved based on CAGE samples from H1 cells with FANTOM ids CNhs14067, CNhs14068 and CNhs13964. All reads with mapping quality less than 10 were discarded. The positive set consisted of CAGE enriched regions (tag-clusters) derived from the +/− 500bp region centered on annotated protein-coding gene TSSs. The negative set was populated with tag-clusters which do not overlap with H3K4me3 and Polymerase II ChIP-Seq peaks and are located in intergenic and exonic/intronic protein-coding gene regions (Figure 1A). For the evaluation process we used CAGE samples from H9 and K562 cells with FANTOM ids CNhs11917, CNhs12824, CNhs12824, CNhs12458, CNhs12684 and CNhs12786 [2]. CAGE tag start site (CTSS) bed files related to these samples were also downloaded and converted to the appropriate format that RECLU and PARACLU can process.

ChIP-Seq datasets in the aforementioned cell-lines were obtained from the ENCODE repository (specific ENCODE sample ids are shown in the following sections). NCBI RefSeq database [18] has been utilized as the reference annotation of coding and non-coding genes that were incorporated in the process of training the ADAPT-CAGE modules and evaluating all algorithms. For the TFBS-oriented evaluation, the genomic coordinates for TFBSs were downloaded from the ENCODE ‘Txn Factor’ track. This track incorporates a large collection of ChIP-Seq derived TFBSs from 161 TFs in 91 cell types.

The evaluation based on epigenomic data was based on the ChromHMM 15-states annotation obtained from the Human Epigenomics [15] repository. E008_15_coreMarks for H9 and E123_15_coreMarks for K562 were downloaded and converted to GRCh38 compatible coordinates. Additionally, H3K4me3 ChIP-Seq derived peaks with sample ids ENCSR716ZJH and ENCSR000EWA for H9 and K562 cells were used. From the RefSeq annotation we retained only protein coding transcripts that have TSS positions at least 100Kb apart, to avoid overlapping with other genic regions. Subsequently, we defined positive and negative zones around annotated TSSs in a two-fold strategy. In the first one, the positive zone was defined as a +/− 500bp window centered on annotated TSSs and the negative zone as a +/− 50kb window centered on same TSSs excluding the positive zone (Figure 4B). For the second strategy, the positive zone was extended to +/− 1kb (Figure 4C).

### Overview of ADAPT-CAGE

The implementation of ADAPT-CAGE is based on a modular approach layering a series of ML models (Figure 1B). It takes advantage of the difference of structural DNA features and Polymerase II associated motifs between promoter and non-promoter regions. The algorithm initially processes aligned CAGE reads to identify loci/peaks enriched in CAGE signal. The nucleotide exhibiting the highest number of overlapping 5’ tag ends is selected as representative for subsequent analysis. Windows of varying size are extracted, centered on the representative nucleotide and vectorized using various structural feature characteristics. Each vector is forwarded into a uniquely trained SVM model to obtain a single probabilistic value representing its significance. A specifically trained Stochastic Gradient Boosting (SGB) model integrates the output values of all SVM models and emits the combined importance of the structural feature set. The affinity of Polymerase II associated motifs (Supplementary Figure 1B) is assessed through 100bp windows around each CAGE peak representative. Polymerase II motifs is integrated into a separately trained SGB model that provides the combined significance of these DNA patterns. The final step of the pipeline, incorporates the structural and Polymerase II motifs with the normalized expression level of each CAGE tag-cluster representative into a final SGB model that distinguishes between transcription initiation events and noise.

### Feature analysis of training dataset

For the quantitative analysis of structural features the sequence around CAGE tag-cluster representatives was converted into numerical profiles (Supplementary Figure 4). Each di- or tri-nucleotide was assigned a value derived from extensive biochemical studies summarized in [20]. A sliding window was subsequently applied to smooth the raw profiles.

Following previous studies [10–13], we analyzed the profile of structural DNA features to observe the aggregated differences between the positive and negative set of CAGE peaks (see the first section in Materials and Methods for the creation of positives and negatives). As expected, similar patterns were observed for the positive set of CAGE tag-clusters compared to the negatives set. One characteristic example of A-philicity is shown in Supplementary Figure 1. Furthermore, we analyzed the profile of thirteen Polymerase II associated motifs (Supplementary Figure 1D) derived from JASPAR database [21]. These motifs represent various promoter-related elements such as Inr, TATA-box, MTE, GC-Box, CCAAT-Box, DPE, BREu, BREd, DCE-S-I, DCE-S-II, DCE-S-III, XCPE1 and MED1. In this case, we also observed striking differences between the positive and negative set. Supplementary Figure 1B shows the INR motif distribution on different categories of said data.

The importance of each feature was assessed with the feature selection module of Boruta [22] package in R (Supplementary Figure 1 C,D). The importance measure of each feature was obtained as the loss of accuracy of classification caused by the random permutation of attribute values between objects. The importance measure (Y-axis) itself varies due to the usage of different data sets for the Polymerase II and structural features. It also depends on the inherent stochasticity of the random forest classifier as well as the presence of non important attributes in the information system (shadow features) [22,23].

The performance of each structural feature fluctuated depending on the combination of the queried sequence size (window around CAGE tag-cluster representatives) and the smoothing parameter (sliding window size). The Polymerase II motif affinity was calculated using TRAP [15,19] and the weight matrices from the JASPAR database [21,22]. The best performing window surrounding the CAGE tag-cluster representatives was found to be of 100 nucleotides in size (data not shown). Details regarding the models’ training are presented in the following paragraphs.

### ADAPT-CAGE training

The positive and negative set of CAGE peaks (see previous section) was split into four subsets, three of which were used for training and one for testing, based on H1 cell samples. Initially, reads with less than 10 mapping quality were removed. From the initial set of tag-clusters only those exhibiting a normalized expression level of more than 1 tpm were retained for downstream analysis. Tag-clusters located closer than 500bp from annotated TSSs and exhibiting an overlap with H3K4me3 and Polymerase II ChIP-Seq peaks consisted the positive set. Tag-clusters overlapping intronic and exonic regions or located in intergenic space, but not overlapping H3K4me3 or Polymerase II peaks, consisted the negative set (as explained in the first section in Materials and Methods). The resulting set of 16,573 (7,614 positives and 8,959 negatives) tag-clusters was divided into 4 non-overlapping subsets consisting of three training sets and one test set for evaluating different combinations of ADAPT-CAGE sub-models. The first training set (3,807 positives and 4,480 negatives) was utilized to train thirteen distinct SVM models using libsvm [24], one for each structural DNA feature, as well as for training the SGB model for the Polymerase II associated motifs. Even though SVM models are theoretically resistant to the curse of dimensionality, the ratio of training samples versus sample dimension was approximately 6:1 to avoid any potential overfitting. Additionally, 10-fold cross validation (CV) was applied during the training process to optimize the generalisation properties of the final models. The second training set (1,524 positives and 1,793 negatives) formed the basis for training the SGB model that combines the results of the individual SVM models related to the structural DNA features. The third training set (1,143 positives and 1,344 negatives) was utilized to train the final SGB model that combines the output of the last tier of SGB models plus the normalized expression level in log_10_ scale. The final set that served as our benchmark consisted of 1,140 positives and 1,342 negatives. Supplementary Figure 4 demonstrates the different sets used for ADAPT-CAGE training and benchmarking.

The assessment of the importance of the final structural and promoter feature models, as well as the normalized expression level for the classification task was performed with Boruta R package (Supplementary Figure 1E). The training of all SGB models was performed with caret package in R [23], using 50 iterations for parameter tuning, and 10-fold CV repeated 10 times to estimate performance and avoid overfitting. SGB was compared against typical Random Forests, Boruta Random Forests, SVM, Self-Organizing Maps and Multilayered Perceptrons and was found to be the best performing algorithm (data not shown).

### Clustering CAGE derived TSSs based on histone mark enrichment

In the FANTOM database, there are 65,141 CAGE tag-clusters characterized as true TSSs in H9 cells. To eliminate the possibility of incorporating CAGE tag-clusters located closely to one-another, thus belonging in the same promoter region, peaks with a distance less than 1kb were merged into a unique event, considering strandness. For each merged event, the position/nucleotide with the highest coverage of CAGE tags was chosen as a representative. This resulted in 31,912 unique transcription initiation events. The whole rationale of this study is that when it comes to utilizing CAGE as a TSS identification methodology, it can be quite noisy. This observation has repeatedly been reported in the literature, and in Figures 2A and 2B we are able to visually inspect this issue in two randomly chosen loci. To assess the severity of the problem genome-wide, we thought of using histone mark ChIP-Seq data in H9 cells from ENCODE, and cluster the 31,912 unique TSS event into groups of similar epigenetic states [25]. To this end, the enrichment of H3K4me3 (ENCFF161EGP ENCODE id), H3K4me1 (ENCFF804YUX), H3K9me3 (ENCFF049TFL), H3K27me3 (ENCFF022SFF), H3K36me3 (ENCFF760NOJ) and H3K27ac (ENCFF271XBU) versus Input (ENCFF734TEC) in the +/− 1kb region surrounding CAGE tag-cluster representatives has been calculated with normR [22]. These enrichment results facilitated the k-means clustering of CAGE tag-clusters in 9 groups, following silhouette coefficient analysis (data not shown). The adjusted p-value was used as a proxy of the enrichment of each mark, to cluster CAGE peaks with similar chromatin states. For visualization purposes, the normalized histone mark ChIP-Seq as well as DNase-Seq (ENCFF110SPC) enrichment was also calculated for 10bp non-overlapping bins spanning the +/− 2kb region centered on CAGE tag-cluster representatives. The same analysis was performed after applying ADAPT-CAGE to the set of 31,912 CAGE enriched loci, separately to the positively (17,850 loci grouped into 10 clusters) and negatively (14,062 loci also grouped into 10 clusters) scored regions (Supplementary Figure 2).

### Genomic location-based evaluation

For the genomic location-based evaluation we considered the overlap of each algorithm’s prediction set with subparts of genic loci (introns, exons and splicing sites) as well as promoter regions (Supplementary Figure 5A). In view of the fact that many predictions can overlap more than one type of annotation, we defined a hierarchy of location:prediction assignments. For instance, a position located 100bp downstream from a TSS could belong to a promoter but overlap the first exon at the same time. Another example could be that candidate TSS events might overlap intronic/exonic regions on a cell-type dependent splicing processing. For this reason, the annotation of candidate TSSs was prioritized with the following strategy. Promoter regions (500bp upstream and 500bp downstream of an annotated TSS) had the highest priority, followed by exon/intron junction regions (+/− 50bp around annotated splice sites). Lastly, exons and introns exhibited the lowest priority.

## Contributions

A.H. supervised the study. G.K.G. envisioned and developed the algorithm and performed the unsupervised analysis. N.P. applied all algorithms and performed all comparisons. A.H., G.K.G. and N.P. made the figures and wrote the manuscript.

## Additional information

### Availability

ADAPT-CAGE is an open source computational framework freely accessible in https://drive.google.com/file/d/1qe15Fd_7Ms5kA6wMdrhkpMe85i4nZg5s/view?usp=sharing.

