## Supplementary Figures for "Solving the transcription start site identification problem with ADAPT-CAGE: a Machine Learning algorithm for analysis of CAGE data"

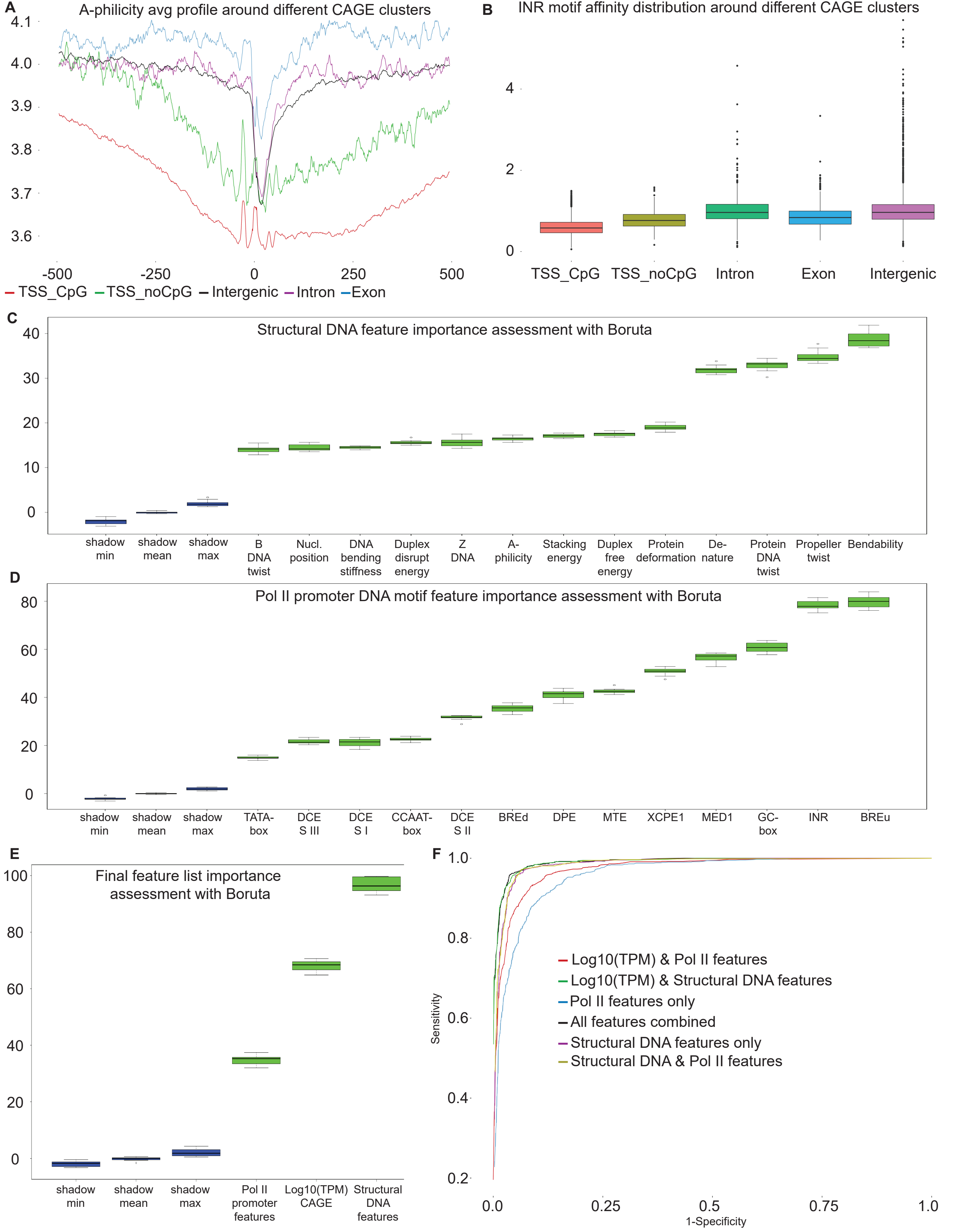
A
A-philicity avg profile around different CAGE clusters
4.1
4.0
3.9
3.8
3.7
3.6
-500
-250
0
250
500
TSS_CpG
TSS_noCpG
Intergenic
Intron
Exon
B
INR motif affinity distribution around different CAGE clusters
4
2
0
TSS_CpG
TSS_noCpG
Intron
Exon
Intergenic
C
Structural DNA feature importance assessment with Boruta
40
30
20
10
0
shadow min
shadow mean
shadow max
B DNA twist
Nucl. position
DNA bending stiffness
Duplex disrupt energy
Z DNA
A-philicity
Stacking energy
Duplex free energy
Protein deformation
De-nature
Protein DNA twist
Propeller twist
Bendability
D
Pol II promoter DNA motif feature importance assessment with Boruta
80
60
40
20
0
shadow min
shadow mean
shadow max
TATA-box
DCE S III
DCE S I
CCAAT-box
DCE S II
BREd
DPE
MTE
XCPE1
MED1
GC-box
INR
BREu
E
Final feature list importance assessment with Boruta
100
80
60
40
20
0
shadow min
shadow mean
shadow max
Pol II promoter features
Log10(TPM) CAGE
Structural DNA features
F
1.0
0.8
0.6
0.4
0.2
Sensitivity
0.0
0.25
0.5
0.75
1.0
1-Specificity
Log10(TPM) & Pol II features
Log10(TPM) & Structural DNA features
Pol II features only
All features combined
Structural DNA features only
Structural DNA & Pol II features

**Supplementary Figure 1. Feature importance assessment and visualization**.

Representative examples of each feature category; structure (i.e. A-philicity) in (A) and promoter-associated motifs (i.e.INR) in (B). The distribution of A-philicity is depicted as an average profile and the distribution of INR affinity as boxplots around CAGE tag-clusters located in different regions of the genome. C) Assessment of each structural feature's importance with the Boruta R package. D) Assessment of each promoter-associated feature's importance. E) Assessment of the importance of the final structural and promoter feature models, as well as the log10(TPM) expression level for the classification task. F) ROC curves of different combinations of models' performance on the test set.

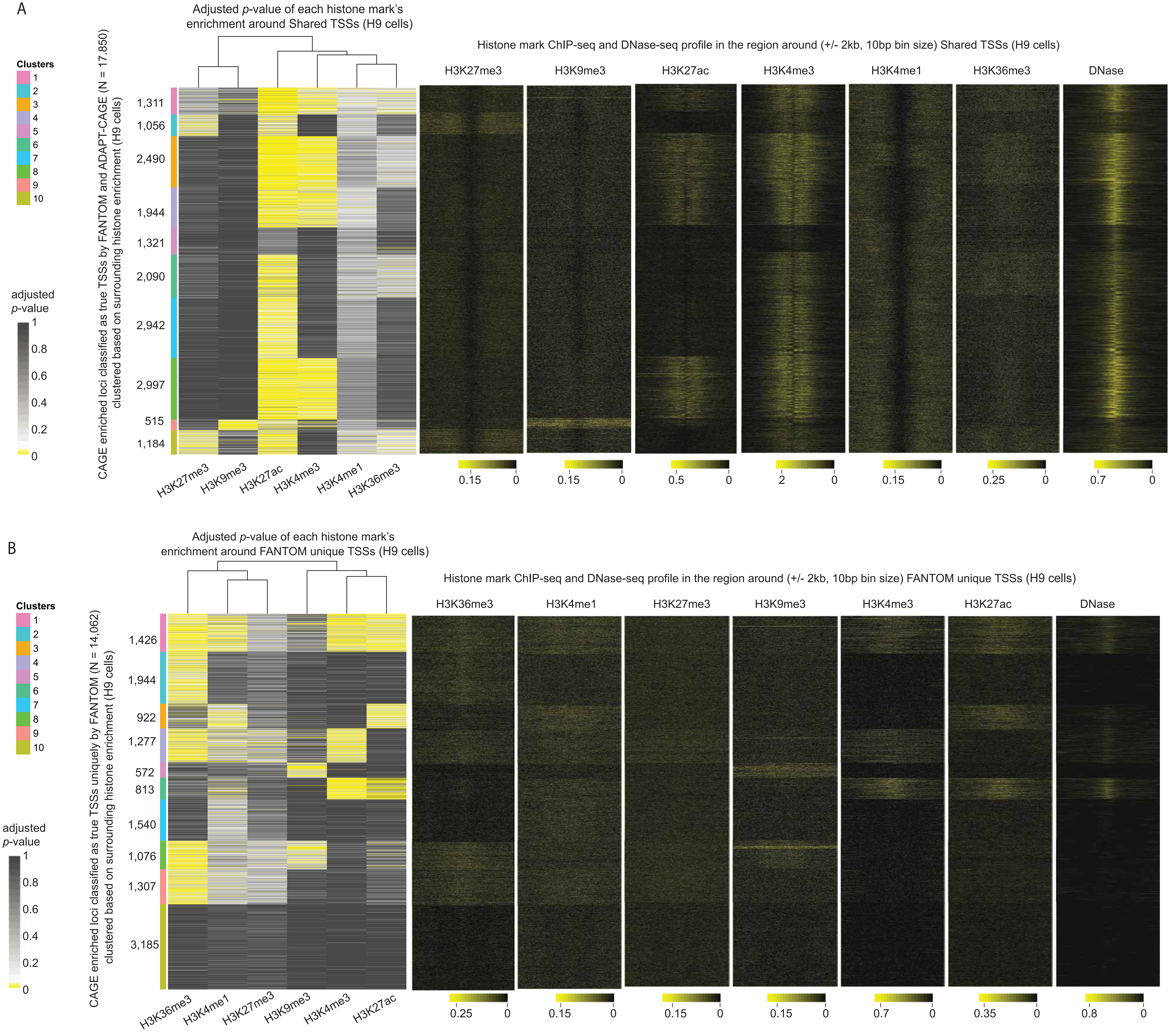
A
Adjusted p-value of each histone mark's enrichment around Shared TSSs (H9 cells)
Histone mark ChIP-seq and DNase-seq profile in the region around (+/- 2kb, 10bp bin size) Shared TSSs (H9 cells)
Clusters
1
2
3
4
5
6
7
8
9
10
adjusted p-value
1
0.8
0.6
0.4
0.2
0
CAGE enriched loci classified as true TSSs by FANTOM and ADAPT-CAGE (N = 17,850) clustered based on surrounding histone enrichment (H9 cells)
1,311
1,056
2,490
1,944
1,321
2,090
2,942
2,997
515
1,184
H3K27me3
H3K9me3
H3K27ac
H3K4me3
H3K4me1
H3K36me3
DNase
0.15
0
0.5
2
0.25
0.7
B
Adjusted p-value of each histone mark's enrichment around FANTOM unique TSSs (H9 cells)
Histone mark ChIP-seq and DNase-seq profile in the region around (+/- 2kb, 10bp bin size) FANTOM unique TSSs (H9 cells)
CAGE enriched loci classified as true TSSs uniquely by FANTOM (N = 14,062) clustered based on surrounding histone enrichment (H9 cells)
1,426
1,944
922
1,277
572
813
1,540
1,076
1,307
3,185
H3K36me3
H3K4me1
H3K27me3
H3K9me3
H3K4me3
H3K27ac
DNase
0.25
0.15
0.7
0.35
0.8

**Supplementary Figure 2. Genome-wide assessment of the differences in the chromatin activity environment surrounding CAGE enriched loci in H9 cells after applying ADAPT-CAGE.**

Clustering of CAGE tag-clusters positively (A) and negatively (B) scored by ADAPT-CAGE, based on the surrounding enrichment of six histone marks' signal. The selected score cutoff was 0.5. The normalized profile of all histone marks' as well as DNase-Seq signal was added as a visual aid.

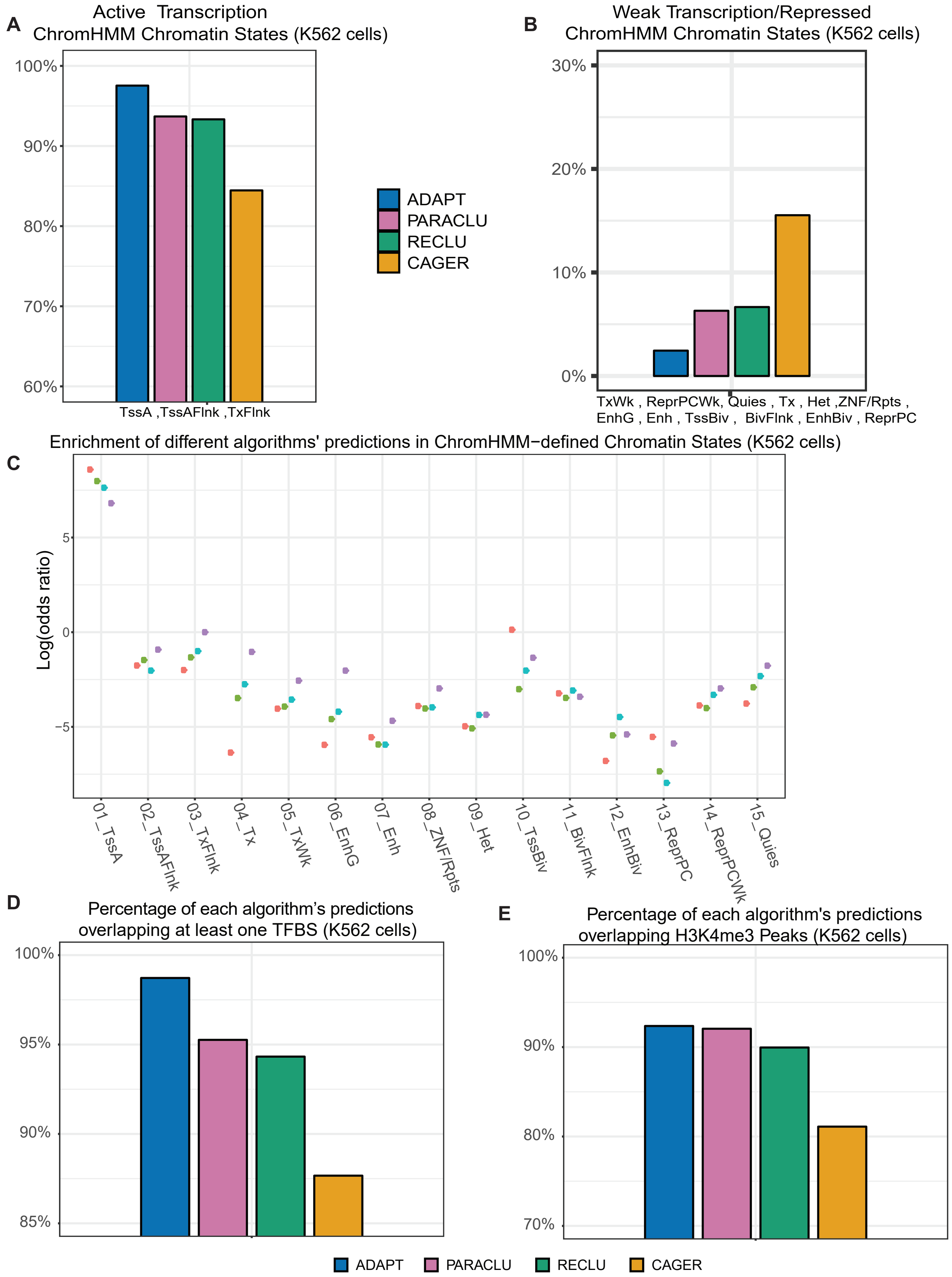
A
Active Transcription
ChromHMM Chromatin States (K562 cells)
100%
90%
80%
70%
60%
TssA ,TssAFlnk ,TxFlnk
ADAPT
PARACLU
RECLU
CAGER
B
Weak Transcription/Repressed
ChromHMM Chromatin States (K562 cells)
30%
20%
10%
0%
TxWk , ReprPCWk, Quies , Tx , Het ,ZNF/Rpts ,
EnhG , Enh , TssBiv , BivFlnk , EnhBiv , ReprPC
C
Enrichment of different algorithms' predictions in ChromHMM−defined Chromatin States (K562 cells)
Log(odds ratio)
5
0
−5
01_TssA
02_TssAFlnk
03_TxFlnk
04_Tx
05_TxWk
06_EnhG
07_Enh
08_ZNF/Rpts
09_Het
10_TssBiv
11_BivFlnk
12_EnhBiv
13_ReprPC
14_ReprPCWk
15_Quies
D
Percentage of each algorithm’s predictions
overlapping at least one TFBS (K562 cells)
100%
95%
90%
85%
E
Percentage of each algorithm's predictions
overlapping H3K4me3 Peaks (K562 cells)
100%
90%
80%
70%
ADAPT
PARACLU
RECLU
CAGER

**Supplementary Figure 3. Evaluating algorithms' performance on experimental data in K562 cells.**

Percentage of each algorithm's predictions uniquely overlapping ChromHMM-derived chromatin states based on the core-15 model in K562 cells. Chromatin states were aggregated in two groups according to levels of chromatin/transcriptional activity. Active (A) and weaker (B) transcription states. C) Enrichment (odds ratio in logarithmic scale) of algorithms' prediction in all ChromHMM-derived chromatin states (TssA and Quies). Percentage of each algorithm's predictions overlapping at least one transcription factor binding site (D) and H3K4me3 (E) ChIP-Seq derived peaks.

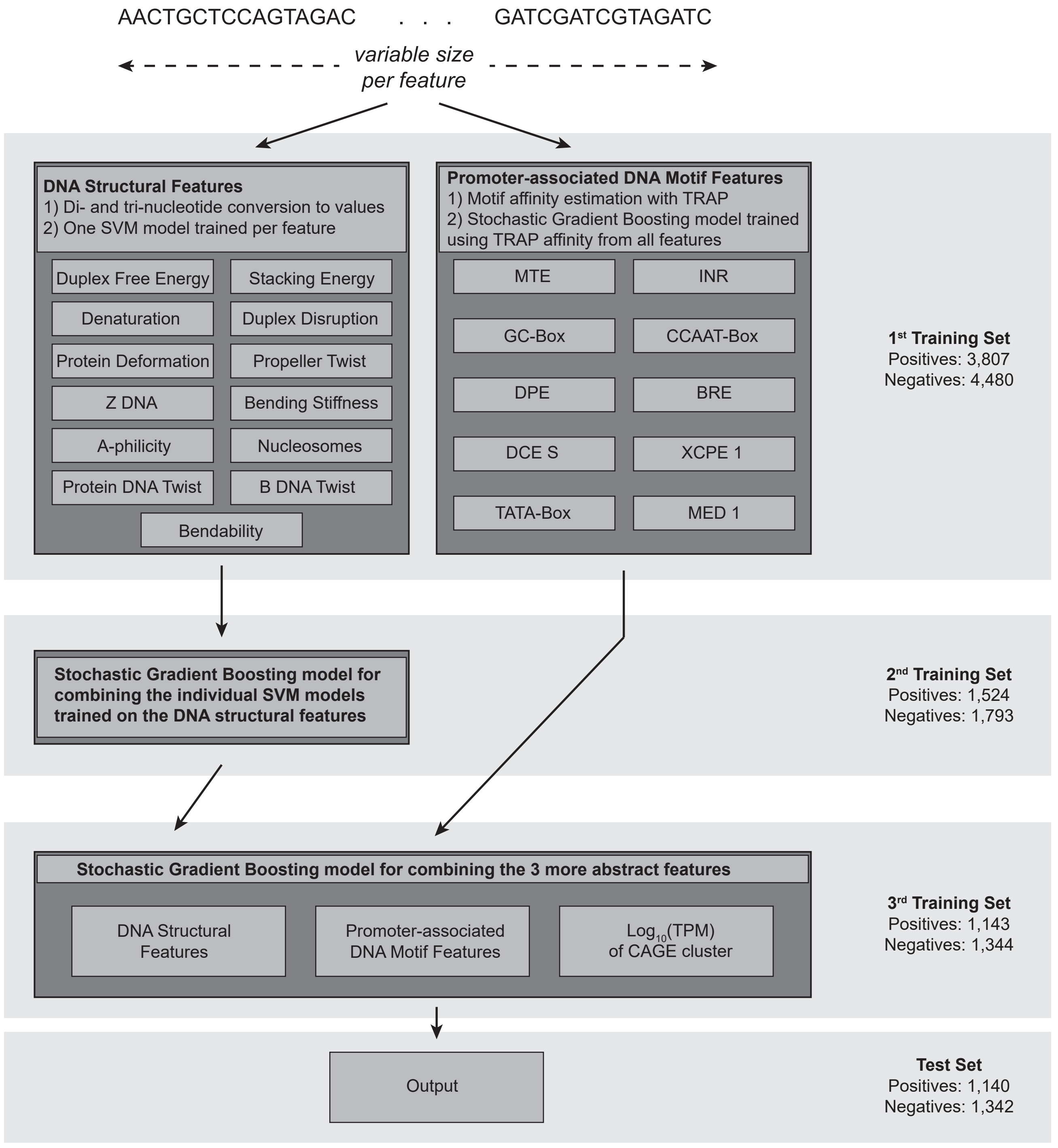
AACTGCTCCAGTAGAC . . . GATCGATCGTAGATC
variable size per feature
DNA Structural Features
1) Di- and tri-nucleotide conversion to values
2) One SVM model trained per feature
Duplex Free Energy
Stacking Energy
Denaturation
Duplex Disruption
Protein Deformation
Propeller Twist
Z DNA
Bending Stiffness
A-philicity
Nucleosomes
Protein DNA Twist
B DNA Twist
Bendability
Promoter-associated DNA Motif Features
1) Motif affinity estimation with TRAP
2) Stochastic Gradient Boosting model trained using TRAP affinity from all features
MTE
INR
GC-Box
CCAAT-Box
DPE
BRE
DCE S
XCPE 1
TATA-Box
MED 1
1st Training Set
Positives: 3,807
Negatives: 4,480
Stochastic Gradient Boosting model for combining the individual SVM models trained on the DNA structural features
2nd Training Set
Positives: 1,524
Negatives: 1,793
Stochastic Gradient Boosting model for combining the 3 more abstract features
DNA Structural Features
Promoter-associated DNA Motif Features
Log10(TPM) of CAGE cluster
3rd Training Set
Positives: 1,143
Negatives: 1,344
Output
Test Set
Positives: 1,140
Negatives: 1,342

**Supplementary Figure 4. Overview of ADAPT-CAGE training process based on CAGE samples from H1 cells.**

CAGE tag-clusters with less than 1 TPM expression level were removed. From the remaining regions, tag-clusters overlapping H3K4me3 and Polymerase II ChIP-Seq derived peaks, located on annotated promoters consisted the positive set. Tag-clusters overlapping intronic and exonic regions or located in intergenic space, but not overlapping H3K4me3 or Polymerase II peaks, consisted the negative set. Both negative and positive sets were subsequently split into 4 sets, 3 of which were kept for training the different layers of the model and 1 for testing the final performance. Raw sequences were scanned to calculate the distribution of each structural feature as well as the affinity of every promoter associated JASPAR motif. Each structural feature was responsible for training its own SVM model while the promoter motifs were combined into an SGB model using the 1st training set. The second training set was used to train the SGB that aggregates the structural DNA SVM models, while the third one was used to train the SGB model that combines the structural and promoter motif features with the log10(TPM) value of CAGE tag-clusters.

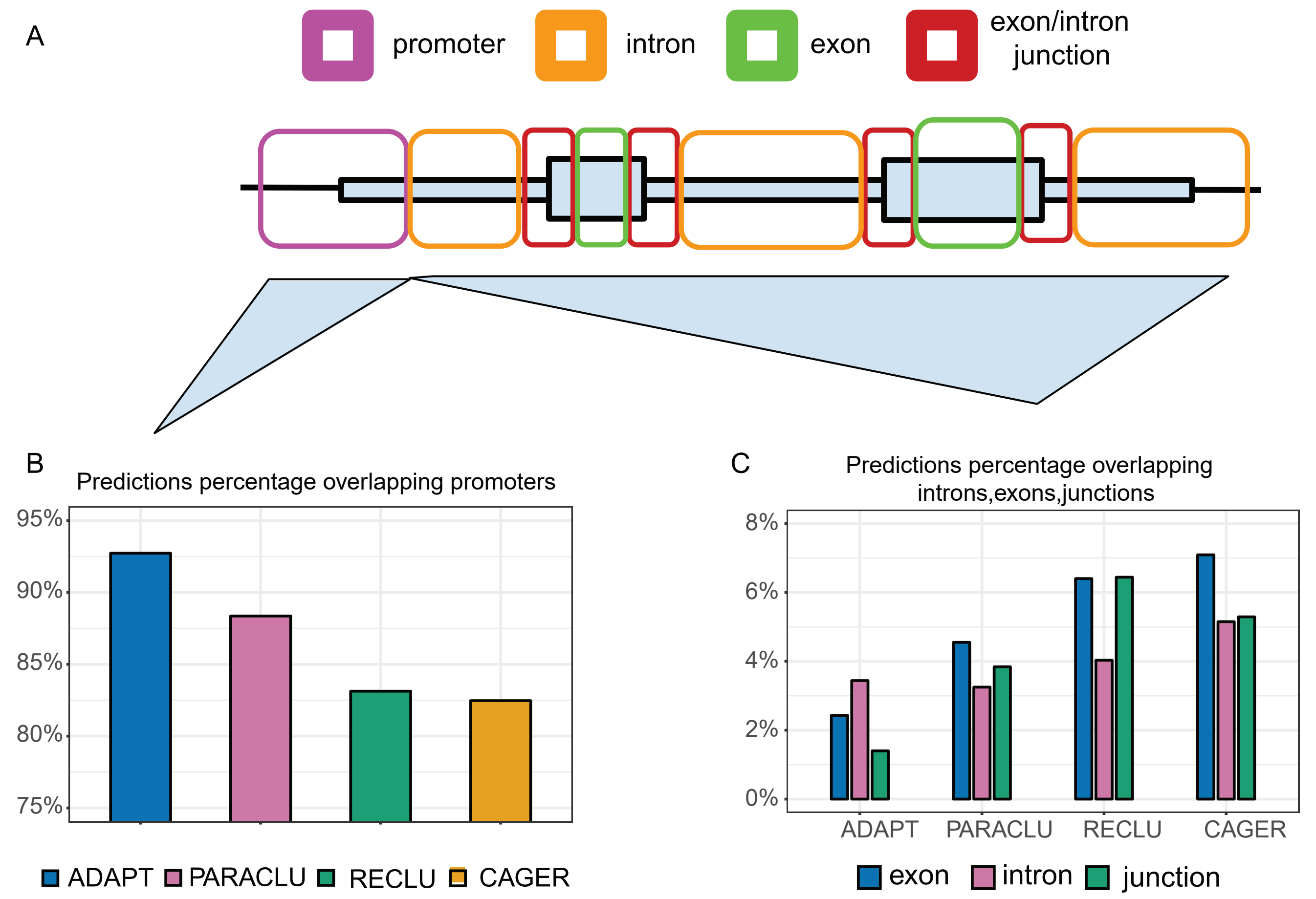
A
promoter
intron
exon
exon/intron junction
B
Predictions percentage overlapping promoters
95%
90%
85%
80%
75%
ADAPT
PARACLU
RECLU
CAGER
C
Predictions percentage overlapping introns,exons,junctions
8%
6%
4%
2%
0%
ADAPT
PARACLU
RECLU
CAGER
exon
intron
junction

**Supplementary Figure 5. Comparison of algorithms' performance based on segmentation of genic regions.**

A) Segmentation of genic regions into different types of annotation such as promoters, introns, exons and exon/intron junctions. Percentage of algorithms' prediction overlapping promoters (B) and other genic types of annotation (C).
