## Supplementary table 1 for "Solving the transcription start site identification problem with ADAPT-CAGE: a Machine Learning algorithm for analysis of CAGE data"

| H9 cell line .CTSS (n= 1.249.782) | | |
|---|---|---|
| K562 cell line CTSS (n=966144) | | |
| algorithm | predictions | |
| | H9 cells | K562 cells |
| Number of CTSS | 1.249.782 | 966,144 |
| FANTOM clusters | 65,141 | 47377 |
| aDaPT (0.5) | 43,169 | 31177 |
| aDaPT (0.9) | 32,268 | 25741 |
| CAGER | 14,948 | 14465 |
| RECLU | 14,749 | 11558 |
| PARACLU | 12,948 | 9453 |

**Supplementary Table 1.** Total number of predictions for each cell line. The first replicate of each experiment is selected for analysis. For ADAPT-CAGE numbers for two different threshold values are listed. For all other algorithms numbers for default parameters are presented (see Supplementary table 4 for default values)
