## Supplementary table 2 for "Solving the transcription start site identification problem with ADAPT-CAGE: a Machine Learning algorithm for analysis of CAGE data"

| State No | MNEMONIC | DESCRIPTION |
|---|---|---|
| 1 | TssA | Active TSS |
| 2 | TssAFlnk | Flanking Active TSS |
| 3 | TxFlnk | Transcr. at gene 5' and 3' |
| 4 | Tx | StronTssAg transcription |
| 5 | TxWk | Weak transcription |
| 6 | EnhG | Genic enhancers |
| 7 | Enh | Enhancers |
| 8 | ZNF/Rpts | ZNF genes & repeats |
| 9 | Het | Heterochromatin |
| 10 | TssBiv | Bivalent/Poised TSS |
| 11 | BivFlnk | Flanking Bivalent TSS/Enh |
| 12 | EnhBiv | Bivalent Enhancer |
| 13 | ReprPC | Repressed PolyComb |
| 14 | ReprPCWk | Weak Repressed PolyComb |
| 15 | Quies | Quiescent/Low |

**Supplementary Table 2.** List of fifteen states model. List of Chromatin states as defined in Human Epigenome Roadmap Project [16] List of fifteen states model. The highly active transcription states are highlighted with green color and these characterized by weak/no transcription with orange color. The grouping based on the hierarchy presented on the [16].
