## Supplementary table 3 for "Solving the transcription start site identification problem with ADAPT-CAGE: a Machine Learning algorithm for analysis of CAGE data"

| number of CTSS for H9 ESC cells (n= 1.249.782) | | | | | | | |
|---|---|---|---|---|---|---|---|
| Fantom Clusters (n=65.141) | | | | | | | |
| algorithm | Cluster Of TSS (p | Lay in | | Overlapping with | | | |
| | | a | b | c | d | e | f |
| | | Near TSS region | [illegible] 50.000) | Histone marks Ac | Histone marks we | TFBS | H3K4me3 |
| FANTOM | 65,141 | | | | | | |
| aDaPT (0.5) | 43,169 | 9456 | 2500 | | | | |
| aDaPT (0.9) | 32,268 | 7740 | 1674 | 30617 | 1667 | 30918 | 29520 |
| CAGER | 14,948 | 3001 | 842 | 12650 | 2353 | 13234 | 12417 |
| PARACLU | 12,948 | 2906 | 582 | 11780 | 1332 | 12067 | 11587 |
| RECLU | 14,749 | 3001 | 842 | 12624 | 2258 | 13076 | 12315 |

**Supplementary Table 3.** Numbers of prediction for evaluations Summary Numbers for annotation (a,b columns) and experimental driven (c,d,e,f columns) comparisons. Each cell contains the quantity of predictions that overlap the column-specific region.
