## Supplementary table 4 for "Solving the transcription start site identification problem with ADAPT-CAGE: a Machine Learning algorithm for analysis of CAGE data"

| Algorithm | Default values | range of testing values |
| --- | --- | --- |
| RECLU | density=2 | desnity (2,4,8,16,32,300) |
| PARACLU | IDR=0.1 | IRD (0.1,0.5, 1, 5, 10 , 30 ) |
| CAGER | Max distance=20 | maxDist (1, 10, 20) |

**Supplementary Table 4.** Algorithm Parameters values used in comparisons. Default values are used unless otherwise stated.
